# RNA Stores Tau Reversibly in Complex Coacervates

**DOI:** 10.1101/111245

**Authors:** Xuemei Zhang, Yanxian Lin, Neil A. Eschmann, Hongjun Zhou, Jennifer Rauch, Israel Hernandez, Elmer Guzman, Kenneth S. Kosik, Songi Han

**Author notes:** **Competing interests** The authors declare no competing financial interests. **Contributions** K.S.K. and S.H. designed the experiments; X.Z. did the PAR-iCLIP experiments and the *in vitro* RNA-protein binding assays; E.G. sequenced the DNA libraries; H.Z. performed the bioinformatics analysis; J.R. did the ITC experiments; N.A.E., Y.L. and J.R. did the RNA-tau *in vitro* droplet formation; N.A.E. did the EPR and DEER experiments; I.H. did the sarkosyl fractionation; Y.L. and H.Z. contributed to the manuscript preparation; X.Z., K.S.K. and S.H. wrote the manuscript.

## Abstract

Non-membrane-bound organelles that behave like liquid droplets are widespread among eukaryotic cells. Their dysregulation appears to be a critical step in several neurodegenerative conditions. Here we report that tau protein, the primary constituent of Alzheimer neurofibrillary tangles, can form liquid droplets and therefore has the necessary biophysical properties to undergo liquid-liquid phase separation (LLPS) in cells. Consonant with the factors that induce LLPS, tau is an intrinsically disordered protein that complexes with RNA to form droplets. Uniquely, the pool of RNAs to which tau binds in living cells are tRNAs. This phase state of tau is held in an approximately 1:1 charge balance across the protein and the nucleic acid constituents, and can thus be maximal at different RNA:tau mass ratios depending on the biopolymer constituents involved. This feature is characteristic of complex coacervation. We furthermore show that the LLPS process is directly and sensitively tuned by salt concentration and temperature, implying it is modulated by both electrostatic interactions between the involved protein and nucleic acid constituents, as well as net changes in entropy. Despite the high protein concentration within the complex coacervate phase, tau is locally freely tumbling and capable of diffusing through the droplet interior. In fact, tau in the condensed phase state does not reveal any immediate changes in local protein packing, local conformations and local protein dynamics from that of tau in the dilute solution state. In contrast, the population of aggregation-prone tau as induced by the complexation with heparin is accompanied by large changes in local tau conformations and irreversible aggregation. However, prolonged residency within the droplet state eventually results in the emergence of detectable β-sheet structures according to thioflavin-T assay. These findings suggest that the droplet state can incubate tau and pre-dispose the protein toward the formation of insoluble fibrils.

## Introduction

Inclusions consisting of the tau protein occur in many neurological conditions with Alzheimer’s disease the most prominent among them. Normally, tau is in a dynamic equilibrium between a microtubule-bound and free state. Under disease conditions tau self-assembles into fibrils that eventually lead to highly insoluble polymeric inclusions known as neurofibrillary tangles. The underlying biophysical basis for the transition of tau from a microtubule-associated protein to an insoluble fibril is unknown. However, a clue comes from the observation that polyanions, such as heparin, promote tau fibrillization [1]. Although less effectively, RNA can also induce tau fibrillization [2, 3], and unlike heparin, RNA is present intracellularly, making it accessible to interact with tau.

Our experiments began with the finding that tau can bind RNA in living cells. Interestingly tau-RNA binding showed selectivity for tRNAs. This observation along with the known categorization of tau as intrinsically disordered and its ability to spread from cell to cell in a manner that resembles prions [4, 5] suggested that tau might share additional properties with other RNA-binding proteins involved in neurodegeneration. These proteins include FUS [6-8], TDP-43 [9], C9ORF72 [10, 11], hnRNPA2B1 and hnRNPA1 [12-14], all of which can undergo liquid-liquid phase-separation (LLPS) from the surrounding aqueous medium into droplets *in vitro*. These highly protein-dense structures, also known in the literature as complex coacervates [15, 16], establish a separated liquid phase typically associated with (1) exceptionally high protein concentration [17], (2) tunability with salt concentration and temperature [18], and (3) multivalent electrostatic interactions involving polyelectrolytes, including RNA, single-stranded DNA and intrinsically disordered proteins (IDPs) [19]. A consensus property of a complex coacervate fluid is low interfacial tension that promotes fusion and coating, and is associated with low cohesive energy between hydrated polyelectrolyte complexes and weakly bound water constituents, consistent with high internal fluid dynamics [16, 20]. Complex coacervate chemistry has been implicated in bio-inspired coating, wet adhesion and engulfment [21-23]. RNA-based coacervates are an important organizing principle for biomolecular condensates in cell biology [17, 19, 24-26].

Here we show that tau-RNA complexation can lead to complex coacervation. When multiple tau molecules weakly bind RNA, and overall charge matching is achieved between the polycation, tau, and polyanion, RNA, tau undergoes reversible condensation and liquid-liquid phase separation into micrometer-sized droplets. Remarkably, within this liquid phase-separated state tau maintains high internal segment mobility and a locally compact conformation that protects the core region of tau known as PHF6(*), as found in dilute solution state, despite the molecular crowding associated with coacervation. In contrast, this region experiences full extension that exposes the PHF6(*) region to the solvent and stacks into β-sheets in the presence of a different polyanion, heparin, that induces irreversible fibrillization of tau [27]. The spontaneous and reversible droplet formation suggests that tau is held in a low energy-barrier fluid state between dilute solution and complex coacervate condensate, with the free energy difference toggled by interactions mediated by ions and hydration water. In fact, a systematic study of complex coacervation as a function of temperature verified the process to be entropy-driven that is likely toggled by the release of counterions and/or hydration water that reduces the net excluded volume of the hydrated biopolymer constituents. However, prolonged residence in this phase state begins to induce β-sheet formation, suggesting that the highly condensed phase state of tau can be a precursor to fibril formation.

## Results

### Tau binds RNA in living cells

RNA binding to tau in living cells was examined by PAR-iCLIP (individual-nucleotide resolution cross-linking and immunoprecipitation) using the human tau-specific antibody, HJ 8.5. Human embryonic kidney (HEK) 293T cells expressing wild type full-length human tau (4R2N), mutant tau (P301L-4R2N) or mutant tau fused to CFP (P301L, 4R1N) were cross-linked, and tau immunoprecipitated with the tau antibody (Fig. 1a-b and Fig. S2a-c). The tau constructs and tau mutants used in this study are shown in Fig S1. Wild type human-induced pluripotent stem cell (hiPSC)-derived neurons (Fig 1c), as well as lines harboring P301L tau and a risk variant for progressive supranuclear palsy, A152T were cross-linked and immunoprecipitated (data not shown). A retinoic acid-differentiated neuroblastoma line (SH-SY5Y) that expresses both tau and the short isoform of MAP2 called MAP2c [28] was also cross-linked and immunoprecipitated (Fig. S2d). ^32^P-labeled RNA bands correspond to immunoprecipitated cross-linked tau-RNA complexes (Fig. 1a-c, lanes 2; Fig. S2a, lane 2; Fig. S2d, lane 2), with strong binding of tau to RNA was observed in all these experiments. Tau-RNA complexes were observed regardless of the tau genotype. PAR-iCLIP experiments with varying RNase concentrations did not shift the ^32^P-labeled band, nor change its intensity (Fig. S2b, lanes 2-3). The radioactive RNA band ran close to the tau protein itself, in contrast to that of most known RNA-binding proteins in the literature [29, 30] that run at a range of higher molecular weights. This radioactive tau band was cut from the membrane, and when sized on an RNA gel, ran in the range of 30 to 100 nucleotides (data not shown). This confirms the presence of RNA, and suggests that tau bound predominantly to small RNAs or RNA fragments. Selectivity was further confirmed by showing that MAP2 did not bind RNA despite its proline rich and microtubule binding domains that are highly homologous to that of tau (Fig. S2d, lane 3).

**Fig. 1:**
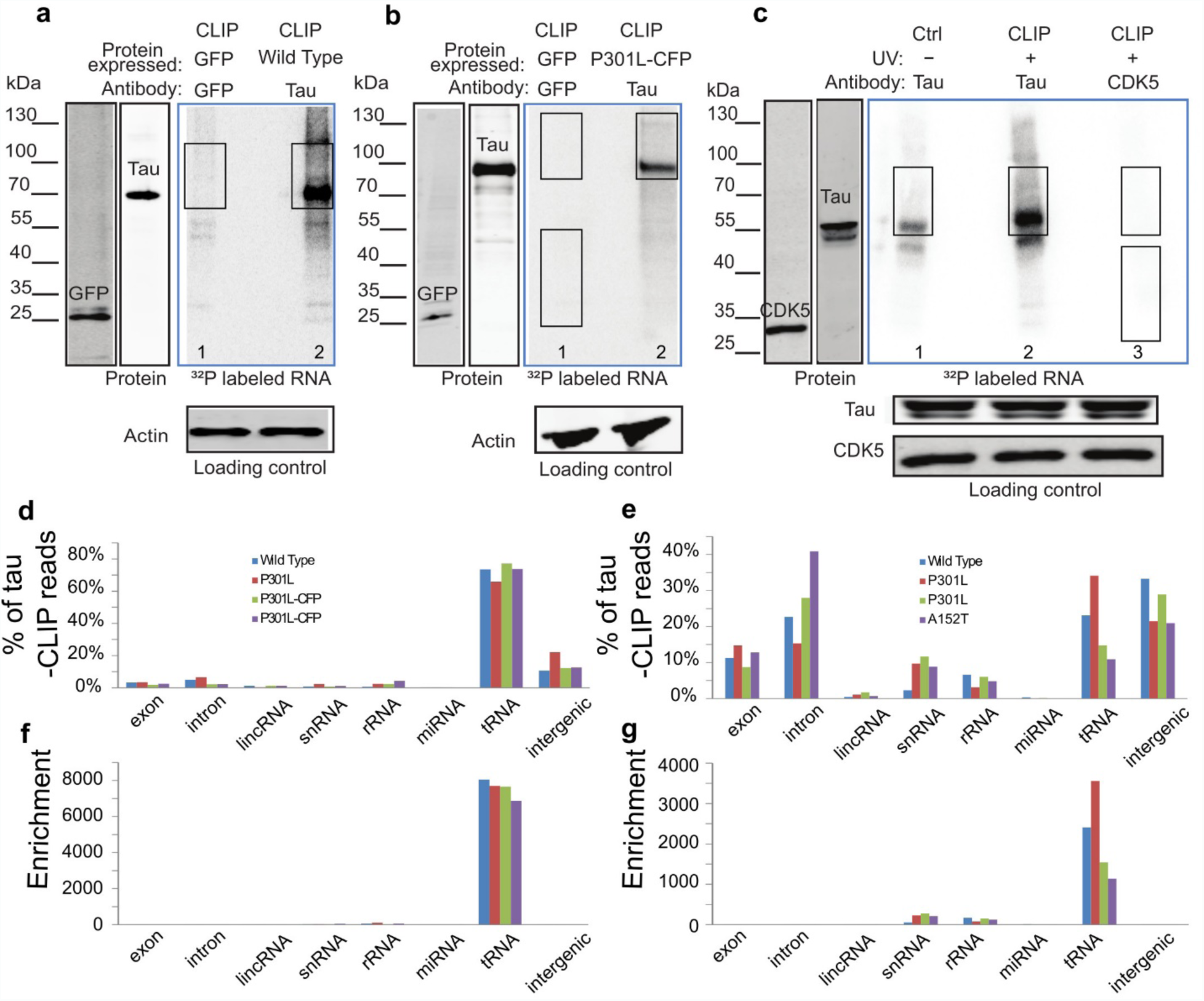
Tau PAR-iCLIP in tau expressing HEK cells and hiPSC-derived neurons. Phosphor images in the blue frames (a-c) show ^32^P-labelled RNA crosslinked to tau protein in HEK cells expressing tau (a-b) and in hiPSC-derived neurons with endogenous tau (c). PAR-iCLIP in HEK cells expressing wild type tau (a, lane 2) or tau P301L-CFP (b, lane 2; *nota bene*: the fused CFP retards the migration of tau). (c) PAR-iCLIP of endogenous wild type tau in hiPSC-derived neurons (lane 2 in c). The antibodies anti-tau HJ 8.5, anti-GFP and anti-CDK5 were used for protein precipitation. No RNase was added, unless specified. PAR-iCLIP with GFP as a control (lane 1 in a-b), with CDK5 as a control (lane 3 in c) and no UV control (lane 1 in c). A small RNA signal was visible in the absence of cross-linking (no UV control) in hiPSC-derived neurons (lane 1, c), suggesting a small portion of RNA may associate with tau *in vitro* after cell lysis. The RNA-protein complexes from CLIP marked by rectangles were cut from the blot for DNA library preparation. Note that two regions of GFP and CDK5 were cut out as sequencing controls in which the lower MW band corresponds to GFP or CDK5. (d-e) % of tau-CLIP reads that are mapped to eight human genome regions in HEK cells (d) and hiPSC-derived neurons (e). (f-g) Enrichment of tRNA in tau-CLIP of HEK cells (f) and hiPSC-derived neurons (g) as discussed in text.

Because small RNAs are abundant and may engage in non-specific interactions, we performed multiple confirmatory controls, including immunoprecipitation in the absence of tau expression, the use of non-tau antibodies rather than HJ 8.5 to rule out non-specific binding, and without UV exposure to eliminate the possibility that tau-RNA complexes formed *in vitro* after cell lysis (Fig. 1a, b lanes 1; Fig. 1c lanes 1, 3; Fig. S2b lanes 1, 4, S2c lanes 3, 4, and Fig. S2d, lanes 1, 3, and 4).

To identify the types of RNA crosslinked to tau, DNA libraries were prepared from the immunoprecipitated radiolabeled bands and sequenced. We analyzed the distribution of tau-bound RNA from the human genome by defining eight regions: exons, introns, lincRNAs, snRNAs, rRNAs, miRNAs, tRNAs and intergenics. In tau-expressing HEK cells, tRNAs were overwhelmingly the highest category of RNA crosslinked to tau (Fig. 1d). Endogenous tau in hiPSC-derived neurons also bound tRNAs; however, background RNA from introns and intergenics were relatively abundant as well (Fig. 1e). Background RNA sequences are common in all CLIP studies, particularly for atypical RNA binding proteins [31]. Despite the abundance of tRNAs in cells, however, background reads from CLIP experiments consistently show a paucity of tRNAs [29, 32-35]. Correcting for background reads by dividing the percentage of tau-bound RNA by the percentage of the nucleotides of each category in the genome demonstrated a selective enrichment of tRNAs that bind to tau in both HEK cells (Fig. 1f) and hiPSC-derived neurons (Fig. 1g). Furthermore, in all experiments, the non-tau controls consistently showed relatively few total reads and very few uniquely aligned reads (Fig. S2e-f).

The specific tRNA species crosslinked to tau overlapped extensively between the HEK and hiPSC-derived neuron samples. Of 625 annotated tRNA loci in the human genome, 462 of the tRNA genes crosslinked to tau in HEK cells, with 79% of them observed in all four tau CLIP samples and 94% in at least two samples. In the hiPSC-derived neurons, all 231 tRNA genes identified were also observed in the HEK cells and 119 of these were verified in at least two tau CLIP samples. The distribution of the tRNAs cross-linked to tau in HEK and hiPSC-derived neurons differed markedly from the total endogenous tRNA distribution. This difference indicated that the tRNAs selected by the CLIP experiment were non-randomly drawn from the total tRNA pool (Fig. S3a-b). Among the most differentially selected tRNAs by CLIP was tRNA^Arg^ (Fig. S3c).

PAR-iCLIP identifies the cross-linked sites in the tRNAs (Fig. 2a-b). We found the tRNA sequences extend from the most 3’ nucleotide of the tRNA to the covalently cross-linked site where the sequencing terminates (Fig. S4). In both HEK cells and hiPSC-derived neurons, the crosslink site was predominantly located within the anti-codon loop followed by the T loop and in the D-loop, but the latter two with far smaller frequencies (Fig. 2b). The observed preference for single-stranded segments of tRNA as crosslink sites to tau, is likely due to the crosslinking between the nucleobases in this region and aromatic rings on the protein. We conclude that tau selectively binds RNA, and the predominant RNA specie bound is tRNA. CLIP procedures are not highly quantitative, and therefore cannot resolve differences in tau-RNA binding among the tau variants tested, P301L or A152T. Therefore, we are unable to conclude whether tau mutations affect its RNA binding.

**Fig. 2:**
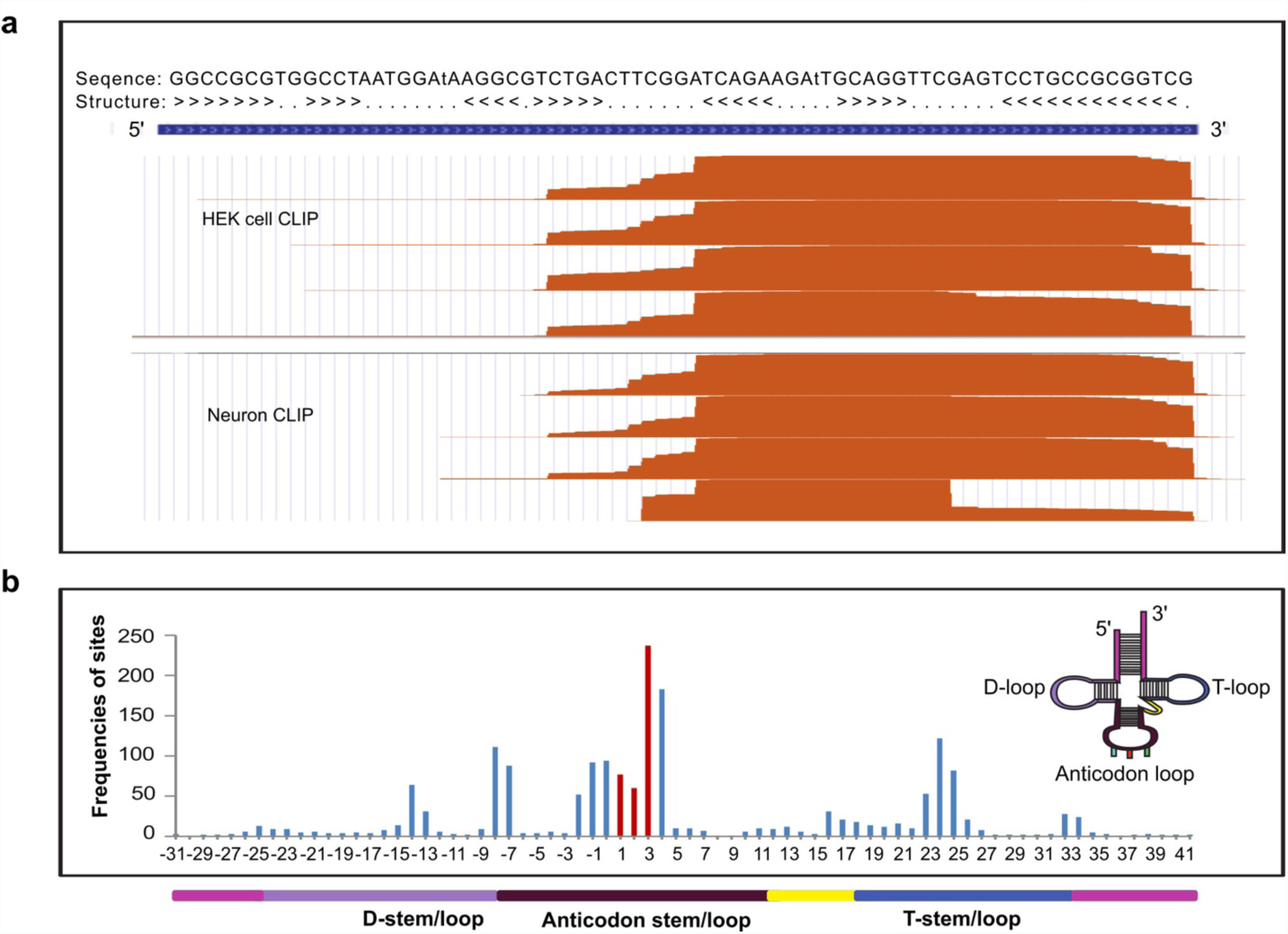
Enrichment of tRNA in PAR-iCLIP data. (a) CLIP cDNA reads from tau expressed in hiPSC-derived neurons (neuron CLIP) and from tau expressed in HEK cells (HEK cell CLIP) that aligned to the chr15.tRNA4-ArgTCG tRNA were found in all CLIP samples, and demonstrated a similar pattern of crosslinking. (b) Analysis of crosslinked sites along tRNA secondary structure demonstrates the anticodon preference, where the anticodon (colored red) region is designated as position 1-3 for alignment purpose. The colored illustration of tRNA secondary structure is displayed as an inset, and below the x-axis in 1-D dimension.

### Tau-RNA binding affinity and stoichiometry

A gel shift assay using recombinant wild type full-length human tau (4R2N) induced a shift in unacetylated tRNA^Lys^ (Fig. 3a), yielding a dissociation constant (K_d_) for 4R2N tau binding to tRNA of 460 ± 47 nM (Fig. 3b). The derived Hill coefficient [36] was 2.8, implying cooperative binding of multiple tau proteins to tRNA. Isothermal titration calorimetry (ITC) experiments independently confirmed the affinity of tau binding to tRNA to yield K_d_ = 735 ± 217 nM and 372 ± 9 nM for 4R2N and K18 tau, respectively (Fig. 3c and Fig. S5a). The dissociation constant for 4R2N binding to a random 43 nucleotide RNA sequence still yielded a K_d_ = 832 ± 94 nM with a Hill coefficient of 2.6 according to a gel shift assay (data not shown), suggesting that tau effectively and non-specifically binds RNA *in vitro*, although there may be differences in binding affinity.

**Fig. 3:**
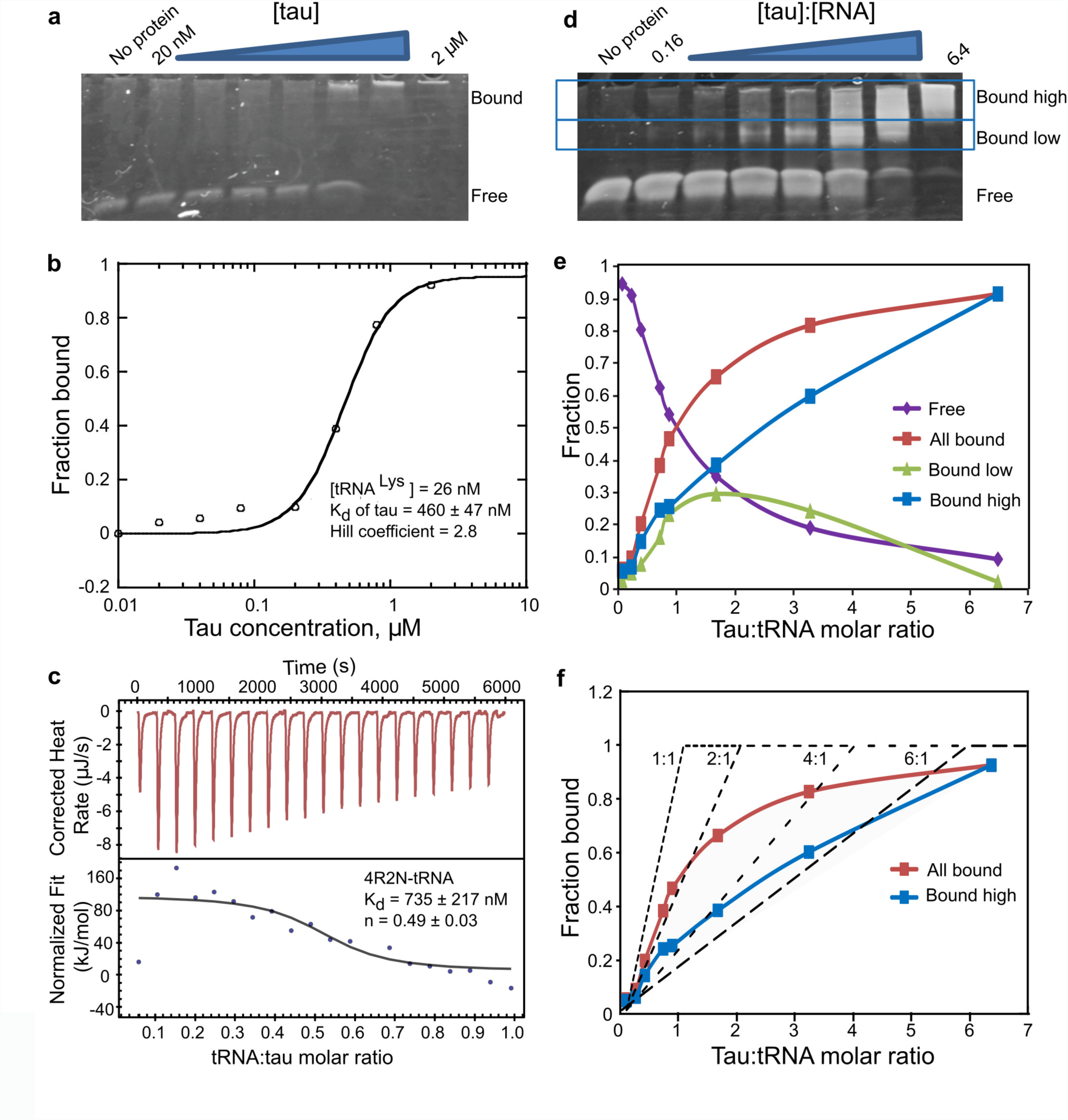
Tau tRNA binding by gel shift assay and ITC. (a) Direct titration experiment shows 4R2N tau induces a mobility shift in tRNA^Lys^. To determine the K_d_ value, direct titration experiment was done, which requires trace RNA concentrations to meet the assumption used in the equation that derives the K_d_ value. The assumption is that the total protein concentration is approximately the free protein concentration at equilibrium, and that protein binding to RNA is negligible. In the direct titration experiments 26 nM tRNA was used and protein concentration spans from 20 nM to 2 μM. (b) The fraction of bound tRNA by 4R2N tau plotted as a function of the monomeric tau concentration and fit to the Hill equation, y = 1 / [1 + (K_d_ / x)^n^]. (c) Yeast tRNA was titrated into solutions of 4R2N tau in an ITC experiment. The top panel in (c) show the raw incremental-titration. The area under each peak is integrated and plotted against the tRNA:tau molar ratio and fitted to an independent binding model (the bottom panel), as discussed in methods. (b-c) Standard error of the mean (SEM) is reported from n=3. (d) The stoichiometric binding experiments were performed by varying the tau:RNA molar ratio at a constant 2.6 uM concentration of tRNA, which approximates the saturation concentration (more than five times the K_d_ of 460 nM). The representative data of three independent experiments is shown. (e) Fraction of bound tRNA from the different bands in (d) is plotted over the molar tau:tRNA ratio. (f) Fraction of bound tau plotted as a function of tau:tRNA molar ratios and compared to the theoretical saturation binding curves (dotted lines) with protein:RNA molar stoichiometries of 1:1 to 6:1. The theoretical curves serve the purpose showing that multiple tau molecules bind tRNA with increasing tau concentrations, while the model is not meant to fit the data, given that multiple populations with different tau:tRNA ratios will coexist.

The gel shift assay showed multiple bands corresponding to different tau:tRNA stoichiometries corresponding to high molecular weight protein-RNA complexes (Fig. 3d, similar gel shift with the tau fragment K18 shown in Fig S5b). The fraction of bound tRNA to 4R2N tau (from the low and high bands) was plotted as a function of tau:tRNA molar ratios (Fig. 3e), and compared to theoretical binding saturation curves representing a range of stoichiometries from 1:1 to 6:1 (see Fig. 3f). The theoretical curves serve the purpose of showing that multiple tau molecules bind tRNA with increasing tau concentrations. The model is not intended for data fitting; rather it shows that multiple populations with different tau:tRNA ratios coexist. However, the data suggests that at lower tau:tRNA molar ratios, tau and RNA predominantly form protein dimer-RNA (P_2_R) complexes, while at higher tau:tRNA molar ratios, tau and tRNA form larger protein multimer-RNA (P_2n_R) complexes with 2*n* reaching as high as 6. This implies that tRNA is capable of binding multiple tau proteins in a multi-step process. Interestingly, when the tau:tRNA ratio was decreased by increasing the tRNA concentration relative to tau, the higher order P_2n_R complexes dissociated to P_2_R, maintaining a tau dimer bound to RNA in the presence of excess (7.5 fold) RNA (Fig. S5c). Such higher stoichiometric signatures were not observed in the ITC measurement, which is less sensitive to binding events associated with small changes in heat. However, ITC titration experiments showed that both 4R2N and K18 tau interacted with tRNA with the stoichiometry of a protein dimer (Fig. S5a n = 0.49 ± 0.03 and Fig. 3c, 0.52 ± 0.04). We conclude that the formation of P_2n_R complexes with *2n* exceeding 2 must be relying on weak interactions between multiple tau proteins and tRNA. A two stage model, in which a protein dimer-RNA (P_2_R) complex forms first followed by the formation of a protein multimer-RNA (P_2n_R) complex, is in fact consistent with the model for RNA binding to the protein AUF1 [37] and hnRNP A1 [38].

### Tau phase-separates in the presence of RNA

Mixing of 4R2N or Δtau187 (similar to K18, see Fig. S1) with tRNA (25 kDa), poly(A) RNA (66∼660 kDa) or poly(U) RNA (800∼1000 kDa) reliably produced a turbid solution under a wide range of tau:RNA mass ratios and salt concentrations. According to bright-field microscopy (Fig. 4a), droplets formed and phase separated from the bulk aqueous phase with a clearly visible and highly spherical boundary. Tau droplets were capable of merging into a single droplet with the complete and nearly instantaneous loss of any boundary at the fusion interface, indicating that the droplets are fluidic with a relatively low interfacial tension (a series of snapshots capturing the fusion of two droplets are shown in Fig. 4a). Confocal microscopy images of fluorescence-labeled tau verified that tau was predominantly contained within the droplet (Fig. 4b). Depending on the specific condition, we were able to observe droplet formation with total Δtau187 protein concentration ranging between 2-400 μM, as will be discussed in greater detail next.

**Fig. 4:**
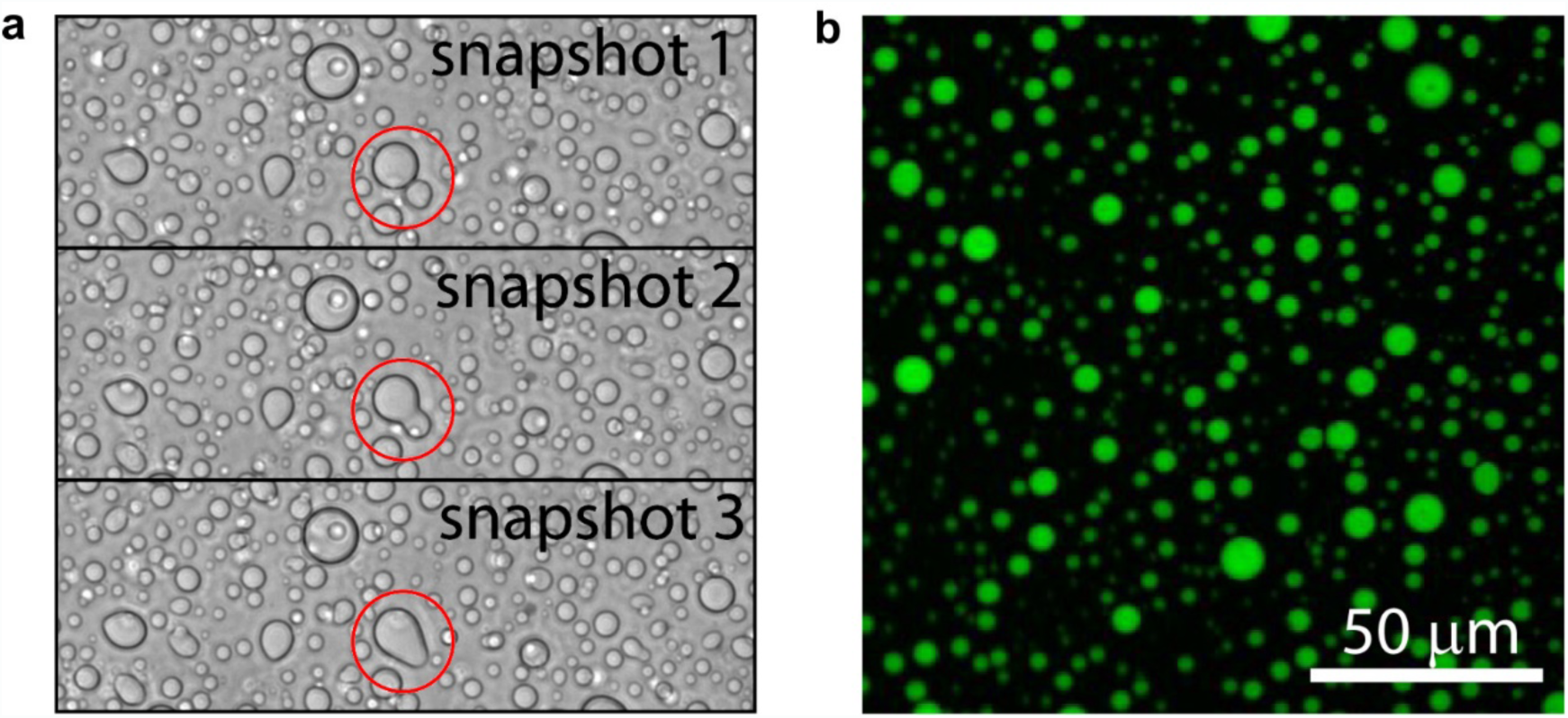
Tau and RNA forms droplet *in vitro*. (a) Bright field snapshots of droplets from 400 μM Δtau187 and 800 μg/ml poly(A) showing two droplets seamlessly fusing (highlighted with red circle). (b) Confocal microscopy image of Δtau187 labeled with Alexa-488 of the droplets from the same sample as in (a). 50 μM Alexa-488 labeled Δtau187 mixed with 350 μM MTSL labeled Δtau187 of droplets formed immediately after adding 800 μg/ml poly(A) RNA. Both (a) and (b) were sampled at room temperature and without added NaCl.

### Tau-RNA droplets form a complex coacervate phase

The amount of tau-RNA droplet was quantified as % coverage from the microscopy images, as well as verified against turbidity measurements relying on light scattering at λ = 500 nm. Droplet formation follows spontaneously from the mixing of two oppositely charged biopolymers, tau and RNA, at a given range of pH, salt and protein concentration, as well as temperature (Fig. 5). This process is fully reversible and reproducible, characteristic of equilibrium states. We highlight the effect of salt concentration first. At a given total protein concentration and temperature, the systematic increase in salt concentration decreased the amount of tau-RNA droplet. Specifically, at room temperature, a total protein concentration of 80 μM and in the presence of tRNA, the droplets reproducibly disappeared when the salt concentration was increased to 100 mM or higher (see Fig. 5a). This general trend was observed repeatedly under a range of experimental conditions, including the use of different RNA types (Fig. 5b, Fig. S6a left panel). Interestingly, droplets formed in the presence of poly(A) and poly(U) tolerated higher than 100 mM salt concentration compared to tRNA. The amount of tau-RNA droplet was also found to be sensitively and reversibly tunable by temperature (Fig. 5d). A sharp dependence of droplet formation on temperature can be seen in the series of images shown in Fig. 5d, where droplets dissolve below 22 °C and appear at and above 23 °C. The observation of a clear increase in the amount of droplet with increasing temperature signifies an entropy-driven process. Consequently, when the temperature was increased to physiological conditions (37 °C), droplet formation was observed at protein concentrations as low as 2.5-5 μM, in the presence of salt concentrations as high as 100 mM and glycerol added as a crowding reagent to mimic the intracellular environment (Fig. S6b). This observation is illustrated in a series of images presented in Fig. 5e—droplets are observed at NaCl concentration as high as 100 mM and tau concentrations as low as 2.5 μM when the temperature is elevated to 37°C, while maintaining the tau:RNA molar ratios. The quantity of droplets decreased with decreasing total tau concentration, and disappeared when tau concentration dropped from 2.5 to 1 μM (Fig. 5e, panel iv). Given that the intracellular concentration of tau is approximately 2-4 μM in neurons [39, 40], conditions under which droplets were observed correspond to protein concentration, salt concentration and temperature resembling physiological conditions.

**Fig. 5:**
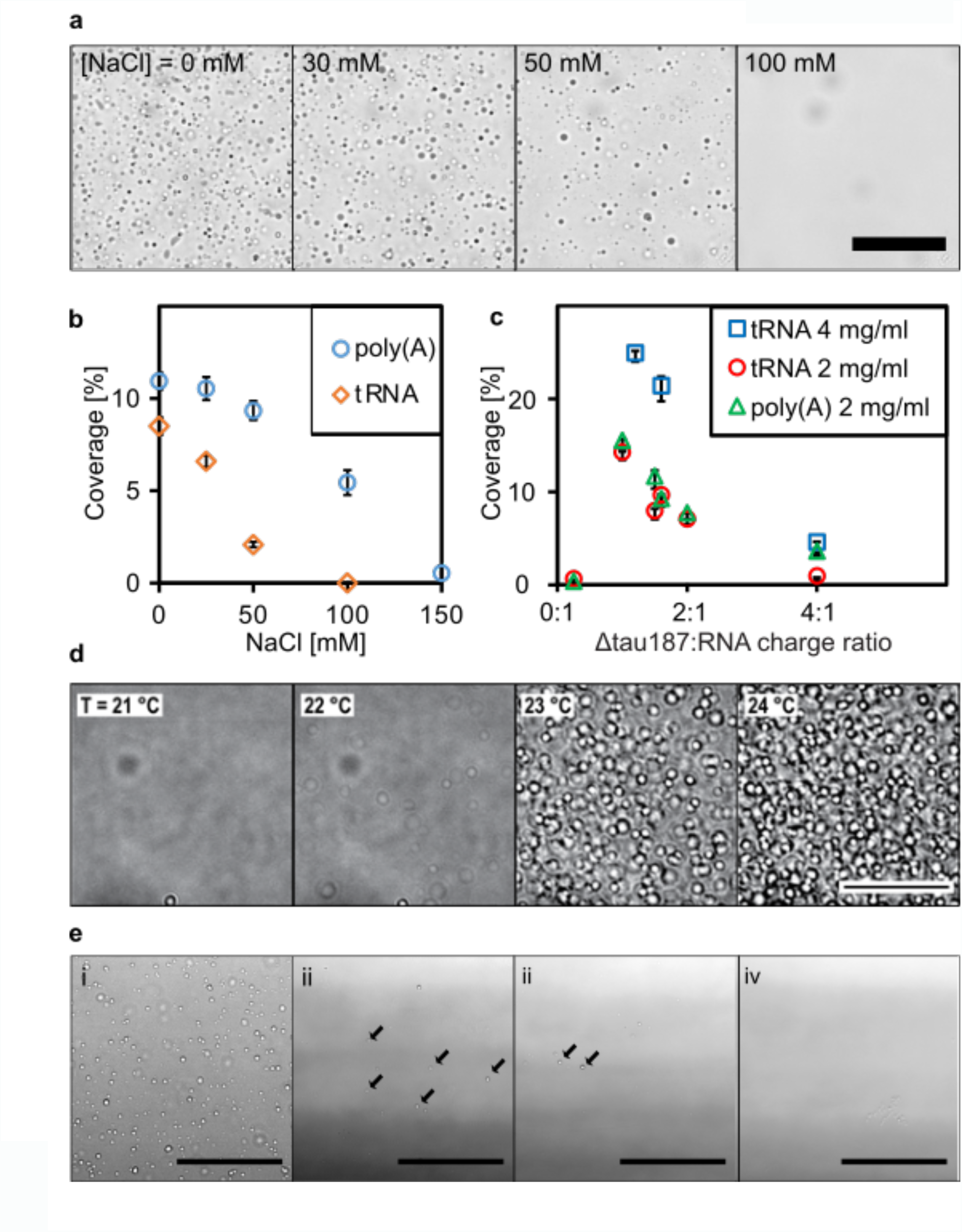
Tau-RNA droplets form a complex coacervate phase. (a) Representative bright-field images of tau-tRNA droplet samples at room temperature with varying [NaCl]. (b) Droplet coverage (in %) with poly(A) or tRNA in room temperature with varying [NaCl]. Δtau187:RNA in a-b were maintained at a mass ratio of 7:1 (corresponding to a charge ratio of 1.2:1) and a total mass concentration of 2 mg/ml. (c) Droplet coverage with poly(A) or tRNA with varying Δtau187: RNA charge ratios. The total mass concentrations are indicated in the legends. Samples made by mixing of 80 μM Δtau187 with 240 μg/ml poly(A)/tRNA or 160 μM Δtau187 with 480 μg/ml tRNA gave the highest droplet coverage (%), which correspond to charge ratio of 1.3-1.2 between tau and RNA. Error bars in (b) and (c) represent the standard deviation from n = 3. (d) Representative bright-field images of tau-RNA droplets as a function of incubation temperature. To record these images, the temperature was ramped from 19 to 25°C at 1°C/min to acquire confocal images with bright field illumination. The samples for these images are generated from 100 μM tau mixed with poly(U) at approximately 1:1 charge ratio in the presence of 30 mM NaCl. (e) Representative bright-field images of tau-RNA droplet samples incubated at 37 °C under otherwise different sample conditions. The concentration for tau, poly(U) RNA and NaCl are i) 5 μM, 15 μg/ml, 0 mM; ii) 5 μM, 15 μg/ml, 100 mM; iii) 2.5 μM, 7.5 μg/ml, 100 mM; iv) 1 μM, 3 μg/ml, 100 mM. Arrows highlight some of the droplets in images ii and iii. Through Fig. 5, scale bars are for 50 μm and all samples were prepared with Δtau187/322C in 20 mM ammonium acetate at pH 7.0.

Droplet formation was maximal, as measured by % coverage from the microscopy images, with Δtau187:tRNA molar ratios of 8:1, Δtau187:poly(A) RNA molar ratios from 33:1 to 330:1 and Δtau187:poly(U) RNA ratios from 267:1 to 333:1 (estimated from the molecular weight of RNAs, see Fig. S6c). By computing the charge (see Supporting Information) and the mass (as measured by the molarity) of the RNA and protein components in the droplet samples, we found that these different molar ratios converge to approximately a 1:1 charge ratios regardless of the type of RNA used, including poly(A) and tRNA species (Fig. 5c) and poly(U) (Fig. S6a, right), and that at different total mass concentration of tau and RNA. Droplet formation was also observed with full length tau 4R2N and RNA at room temperature, but was less robust than with Δtau187, possibly due to the additional negative charges at the N-terminus of tau 4R2N that would diminish the electrostatic association between tau and RNA. However, at pH 6 where the net charge of tau 4R2N is similar to Δtau187, droplets reliably formed (Fig. S6d), albeit with a significantly lower yield (Fig. S6e)—note the % coverage with tau 4R2N-derived droplets of order < 3% compared to that with Δtau187-derived droplets of order 5-30%.

The observations that droplet formation is directly, sensitively and reversibly tunable by salt concentration, tau:RNA charge ratios, as well as temperatures demonstrate that this process is modulated and toggled by both electrostatic interactions and changes in net entropy, and the balance between these contributions. Based on these findings, we conclude that tau-RNA droplet formation follows a complex coacervation mechanism, as initiated through non-specific and relatively weak complexation of oppositely charged and polyelectrolytes, and driven by the further association of these polyelectrolyte complexes to form a macroscopic fluid phase.

### Tau in condensed droplets assumes solution state properties

Within a condensed complex coacervate fluid, held together by non-specific and weak electrostatic interactions, we expect the polyelectrolyte constituents to maintain their hydration layer and remain dynamic [20, 41]. However, this assumption needs experimental verification. To spectroscopically track tau exclusively from within the condensed phase, we first verified by confocal fluorescence imaging that Δtau187 was predominantly localized within the droplet (see Fig. 4b). This allowed us to characterize the droplet-internal protein properties using spin labeled Δtau187. We carried out continuous wave electron paramagnetic resonance (cw EPR) spectral line shape analysis of singly spin labeled Δtau187 at a cysteine site 322, Δtau187/322C-SL, diluted with diamagnetically labeled Δtau187/322C-DL (see Methods) to compare the protein side chain dynamics and degrees of freedom of the tethered spin label of Δtau187 in dilute solution state (Fig. 6 red, a-c), in the droplet state associated with poly(A) RNA (blue, a) or tRNA (green, b), and upon addition of the tau aggregation inducer, heparin (black, c). Remarkably, the EPR lineshape of Δtau187/322C-SL within the droplet phase was indistinguishable from that in the dilute solution state (e.g. see red vs blue trace in Fig. 6a). In contrast, the EPR line shape dramatically broadened within minutes of adding heparin—a highly effective aggregation inducer of tau [1, 42] (e.g. see red vs black trace in Fig. 6c). This finding experimentally demonstrates that the condensation of Δtau187 to high protein concentration, as found within droplets, alone is insufficient to induce dehydration and aggregation of tau, and that tau retains the protein dynamical properties as found in solution state, despite forming long-range associations with RNA in a highly concentrated fluid phase. From this, we infer that tau maintains its protein dynamics and hydration shell as in dilute solution state.

**Fig. 6:**
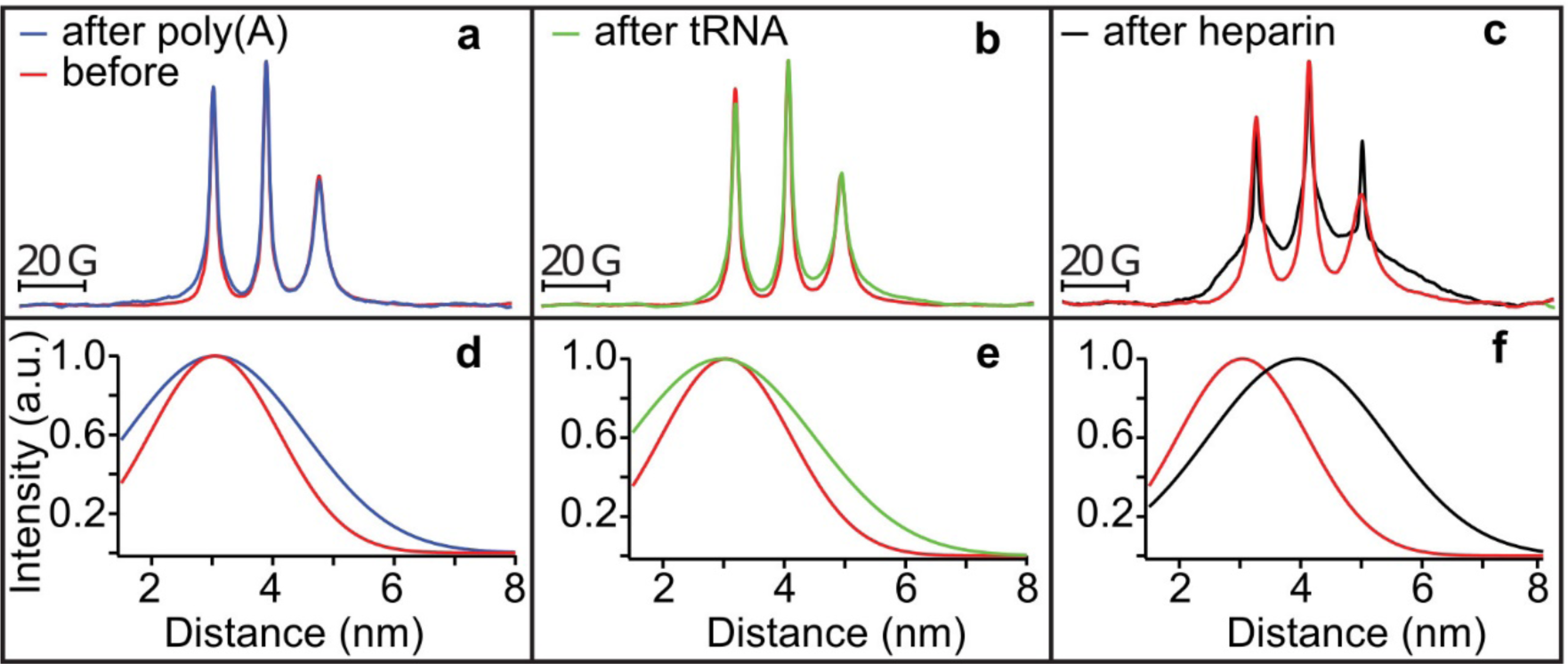
Tau in condensed droplets assumes solution state properties. (a-c) Cw EPR spectra obtained at room temperature of 500 μM Δtau187-SL in droplets formed with 1.5 mg/ml poly(A) RNA (blue in a) and 1.5 mg/ml tRNA (green in b) is unaltered from solution before adding RNA (red in a and b). Cw EPR line shape upon adding 125 μM heparin (black in c) show dramatic line broadening (compared to red in c). DEER of Δtau187-SL_2_ in droplets formed with 1.5 mg/ml poly(A) RNA (blue in d) and 1.5 mg/ml tRNA (green in e), as well as upon incubation with 137.5 μM heparin (black in f) compared to Δtau187-SL_2_ in solution (red in d-f).

To determine whether tau assumes altered protein conformations in the complex coacervate phase compared to the dilute solution phase, we compared the intra-protein distances flanking the PHF6 and PHF6**^*^** hexapeptide regions of tau under these conditions. These regions of the tau sequence pack into the β-sheet core once fibrils are formed of tau [43]. Recently, we identified that both the PHF6 and PHF6**^*^** hexapeptide regions of tau undergo a dramatic opening from a compact to a fully extended local conformation, well before fibril formation is observed, and remain extended as they pack into β-sheet structures as part of insoluble fibrils [27]. These local distance measurements of the regions flanking the PHF6^(^* ^)^ regions offer a powerful tool to compare the conformational state of tau in dilute solution, droplet and aggregation-induced states. We prepared a Δtau187 G272C/S285C-SL_2_ construct that was doubly MTSL-labeled at sites 272 and 285 (see Methods) to access the distances across the PHF6**^*^** hexapeptide region by double electron-electron resonance (DEER) spectroscopy. Surprisingly, the mean distance flanking the PHF6* region remained unchanged from dilute solution state to when tau was condensed into a concentrated complex coacervate phase in association with poly(A) RNA and tRNA (compare blue vs red trace in Fig. 6d, and compare green vs red trace in Fig. 6e). This contrasts with the effect of heparin on tau that markedly extended the mean distance between the labels (from ∼3 to ∼4 nm) within minutes of heparin addition as recently reported [27], corresponding to the extended conformation that the PHF6^(*)^ segment adopts when neatly stacked in β-sheets (compare black vs red trace in Fig. 6f).

Interestingly, we observed a low-level Thioflavin T (ThT) fluorescence under tau-poly(U) RNA droplet forming conditions that gradually increased over 15 hours (see Δtau187 + poly(U)). The ThT fluorescence, however, was nearly eliminated at increased salt concentration that corresponded to conditions under which tau-RNA associations are weakened and droplets are dispersed. However, even after 15 hours of incubation, the ThT fluorescence intensity from the tau-RNA droplet samples was significantly less (less than 5%) compared to what was observed in the presence of the aggregation-inducing heparin, under similar charge ratio and mass concentration and following a brief incubation time of less than 20 minutes (see Fig. S6f, the histogram for Δtau187 + poly(U) vs heparin is not to scale, as indicated with a split y axis). Since ThT fluorescence is commonly used as an assay to detect β-sheet content, we suggest that droplet formation through association with RNA increases the aggregation propensity of tau *in vitro*, even when the tau-RNA complexes are held together by reversible and weak interactions between intact RNA and the hydrated tau constituents.

### Exogenous tRNA can induce sarkosyl insolubility of tau

To determine whether tau-RNA complexes have the potential for pathological interactions *in vivo*, hiPSC-derived neurons with a P301L mutation or wild type were transfected with 48 μg tRNA per 1.2 million cells. The uptake of tRNA was demonstrated with tRNA^Phe^ - fluorescein (Fig. S7). Cell lysates (input) were prepared in a high salt/high sucrose buffer, followed by fractionation in a 1% sarkosyl buffer. Transfection in the absence of nucleic acids (mock) or addition of tRNA to the mock lysate (mock + tRNA) were used as controls. Cells transfected with tRNA accumulated sarkosyl-insoluble tau as detected with the PHF-1 tau antibody; whereas cells without added tRNA, or when tRNA was added to the lysis buffer did not increase the tau population in the insoluble fractions (Fig. 7a). The increase in sarkosyl-insoluble tau populations occurred in both P301L mutation and wild-type tau cells (Fig. 7b-c) when infected with tRNA.

**Fig. 7:**
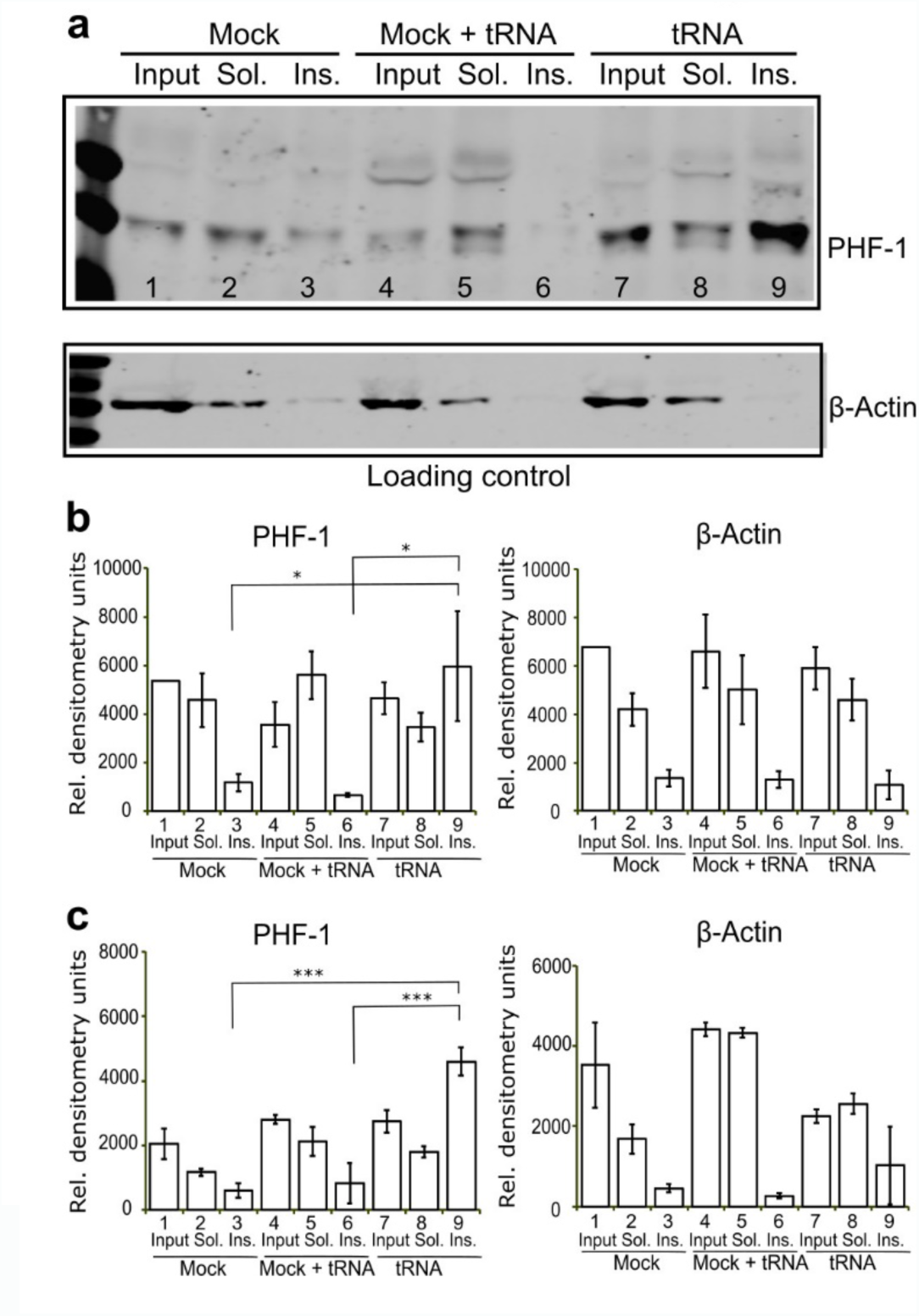
tRNA transfection accumulates sarkosyl insoluble tau in hiPSC derived neurons. (a) Representative western blot of neurons harboring a P301L *tau* mutation transfected with tRNA, but not cells transfected in the absence of nucleic acids (Mock) present evidence for accumulation of sarkosyl insoluble tau (Ins.) as seen in the intense band in lane 9 labeled with PHF-1 antibody. Addition of tRNA to the lysis buffer is insufficient to increase the tau present in the insoluble fractions (compare Mock + tRNA, lane 6 to tRNA, lane 9). (b-c) Quantification of PHF-1 tau and β-actin level for both neurons harboring a P301L tau mutation (b) and neurons expressing wild type tau (c) are shown. Error bar represents standard error of the mean. * p < 0.05, * * * p < 0.001, n=5.

### Discussion and Conclusion

Many years ago Wolde and Frenkel [44] have proposed that condensed liquid droplets serve as a “metastable crucible” to lower free energy barriers for crystal nucleation, while the discovery of the involvement of nucleic acids in coacervation-driven liquid-liquid phase separation (LLPS) dates back to 1949 by Bungenberg de Jong [21]. Here, we identify tau to undergo complex coacervation upon association with RNA. We find that tau can bind RNA in two stages, mediated by (i) strong binding at a tau:RNA molar ratio of 2:1 with nanomolar dissociation constants and (ii) weak association between tau multimers and RNA to form higher order complexes. Although tau lacks a recognizable RNA-binding motif, it selectively bound tRNA as its dominant partner *in vivo*. Furthermore, tau was found to spontaneously phase separate upon non-specific complexation with RNA into dense protein droplets, whose formation was reversibly and sensitively tunable by salt concentrations, tau:RNA ratios, as well as temperature. Crucially, the optimal tau:RNA molar ratios range from 8:1 to 330:1, which varied among different RNA species. Remarkably, all higher order complexes converge to a common tau:RNA charge ratio of ∼1:1. This finding together with the sensitivity of droplet formation to salt concentration led us to conclude that tau-RNA droplet formation is driven by complex coacervation, in which oppositely charged polyelectrolytes associate into extended assemblies held together by electrostatic interactions. Significantly, EPR spectroscopy verified that tau remained locally dynamic, and maintained local conformations as found in solution state. This is consistent with an entropically-driven complex coacervation process between tau and RNA, in which contribution from attractive tau-RNA interactions and changes in the biopolymer conformational entropy are small. We rationalize weak tau-RNA interaction to water-mediated electrostatic attraction between hydrated tau and RNA molecules, that is hence weakened. The total entropy of the complex coacervated system can be increased through the release of counterions and/or hydration water, provided that the entropy of the biopolymer constituents themselves remains relatively constant. That the release of hydration water may be one of the potential drivers of entropically-driven complex coacervation is not contradictory to our statement that fully hydrated biopolymers constitute the complex coacervate phase. Even sharing the hydration shell between proteins in a crowded solution alone will increase the net entropy of the total system’s solvent, if the protein concentration within the coacervation is very high and the protein preserves its dynamics and hydration shell as in dilute solution state. Notably, there are several reports in the literature on the spontaneous complex coacervation of synthetic polyelectrolyte and protein (e.g. β-lactoglobulin) constituents to be entropy-driven [6, 18, 24, 45, 46], but no studies combine macroscopic thermodynamic measurements with studies of local protein dynamics and conformations. We show that the majority of tau complexed with RNA in the highly condensed droplet state maintains locally the dynamical conformations of free tau as populated in dilute solution state. In other words, tau can reversibly switch between the dilute solution and the dense droplet state involving minimal rearrangement of hydration water and protein conformations, as tuned by salt concentration, pH and temperature—i.e. physiologically viable “knobs”.

Still, a low-level ThT fluorescence was observed in the tau-RNA droplet samples after prolonged incubation, but with a significantly reduced amplitude compared to tau incubated with heparin. Consistent with this finding, tRNA increased sarkosyl insolubility of tau *in vivo*. We conclude that dynamic and free tau species are stored in a concentrated droplet state that can spontaneously and reversibly dissolves into solution at increased salt concentration and/or depressed temperature, while also being predisposed to converting to fibrils. This previously unrecognized droplet phase state contrasts with that of tau following incubation with the polyanion, heparin, in which a rapid conformational change occurs in solution state followed by an irreversible fibril formation. The distinguishing effects between RNA (charge density ∼3.0/nm, estimated from the nucleic acid double helix [47], a polyphosphate) and animal-derived heparin (charge density ∼4.0/nm, estimated from a disaccharide crystal structure [48], a polysulfate) on tau—coacervation versus fibrillization—may be an operational principle of tau-polyanion association.

Linking the *in vitro* observations of tau-RNA coacervation to *in vivo* structures that share many of the properties we describe here remains a challenge for the field. RNA granules [49-51], stress granules [52], germ line granules [53], synaptic vesicles [54], chemical reaction centering at the origin of life [55] all lack taxonomy, and yet share features that resemble *in vitro* droplets. These structures are cohesively motile in the absence of any surrounding membrane, contain IDPs which often contain RNA-binding proteins and serve as an RNA transport vehicle in which translation is silenced [51, 56, 57]. TDP43, FUS, C9ORF72, hnRNPA2B1 and hnRNPA1 and TIA-1 [6-14] are all IDPs and RNA binding proteins, but establishing a connection between *in vivo* systems and the biophysics of droplets has been challenging. Brangwynne et al have recently employed optogenetic control of Cry2 to trigger the association between IDPs tethered to Cry2 as a mechanism to induce phase separation involving the IDPs, presumably by increasing the local protein concentration. Our study not only adds tau to this prominent list of proteins that undergo LLPS upon association with RNAs, but also establishes a new phase state of tau in which soluble tau is reversibly stored in concentrated complex coacervate droplets, readily bioavailable for cellular functions or vulnerable to pathological aggregation depending on the environmental cues at hand, and offers guidance for the physiologically tunable factors that may promote LLPS *in vivo* – increased temperature, lower salt concentration, and exceeding a local threshold protein concentration.

## Materials and Methods

### Cell culture

A total of eight samples were used for PAR-iCLIP studies. Four samples were human embryonic kidney (HEK 293T) cells that expressed either 4R2N (residues 1-441) wild type *tau* (one sample), 4R2N *tau* harboring the P301L mutation (one sample), and 4R1N *tau* harboring the P301L mutation fused to CFP (two samples). Four samples were neurons derived from human induced pluripotent stem cells (hiPSCs) obtained by reprogramming dermal fibroblasts with virally transduced Yamanaka factors [58] that expressed wild type *tau*, two harboring a P301L *tau* mutation and one harboring the A152T variant [59, 60]. The hiPSC lines were karyotyped by Cell Line Genetics. Both wild type and P301L were from normal males (46XY). A152T displayed a balanced three-way translocation of chromosomes 1, 13 and 7, which most likely occurred during the reprogramming process (46XY, t(1;13;7) (q31.2;q21;q36.3). Pluripotent cells were maintained under feeder-free conditions and cultured in BD Matrigel (BD Biosciences) coated six-well plates and fed with mTSER daily (Stemcell Technologies). Neuroectoderm differentiation utilized dorsomorphin and SB431542 (Sigma-Aldrich) for a week [61]. When neural-rosettes were clearly observable, the media was gradually replaced with neuronal induction media containing Knockout DMEM F12 with N2, and Glutamax 1X supplemented with laminin (Sigma-Aldrich) and maintained for an additional six days with every other day feeding. Neuro-rosettes were microdissected, and grown as neurospheres in the neuronal induction media supplemented with B27 supplement. Neurons were differentiated from neurospheres by dissociating them into individual cells enzymatically with a 1/1 mixture of 0.25% trypsin / StemPro accutase (Thermo Fisher), and plated on Poly-L-Ornithin/Laminin coated six well plates at a density of 200,000 cells per well. Neuron differentiation and maturation were stimulated by the addition of 10 mM each of NT3, BDNF, and GDNF (Preprotech) to the neuronal growth media containing Neurobasal, N2 supplement, B27 supplement and Glutamax. Neurons were fed twice per week by replacing half of the conditioned media with pre-warmed supplemented fresh media, and underwent maturation for at least five weeks. At this developmental stage, hiPSC-derived neurons predominantly express the 3R tau isoform. Neuroblastoma (SH-SY5Y) cells were plated as monolayers in DMEM/F12 medium with 10% FBS at 1x 10^6^ per 10 cm dish, and the cells switched to neurobasal medium the next day and differentiated for seven days using 10 μM retinoic acid.

### Antibodies

The tau antibodies used were Tau-5 for probing total tau in western blot (1 mg/ml, Abcam); human tau-specific HJ 8.5 and mouse tau-specific HJ 9.2 (4.5 mg/ml and 2.8 mg/ml, both HJ antibodies are gifts from Dave Holtzman, Washington University) for CLIP; PHF-1 (gift from Peter Davis, Albert Einstein College of Medicine) for sarkosyl-insoluble tau. Other antibodies used are MAP2 antibody (0.3 mg/ml, Proteintech^TM^); CDK5 rabbit antibody (0.2 mg/ml, Santa Cruz); GFP rabbit antibody (5 mg/ml, Abcam); β-Actin (Sigma-Aldrich); mouse IgG_1_ Alexa 680, (2 mg/ml, Invitrogen); mouse IgG Alexa 680 (2 mg/ml, Invitrogen); rabbit IgG 800 (1 mg/ml, Odyssey).

### PAR-iCLIP

PAR-iCLIP (<u>P</u>hoto<u>A</u>ctivatable <u>R</u>ibonucleoside-enhanced <u>I</u>ndividual-nucleotide resolution UV <u>C</u>ross-<u>L</u>inking <u>I</u>mmunoPrecipitation and high-throughput sequencing) was applied to detect specific RNAs bound to tau in living cells [30, 32]. Cells were treated with 4-thiouridine (4SU) (Sigma-Aldrich) for 1h at a final concentration of 500 μM at 37°C, rinsed with ice-cold 1x PBS and irradiated one time with 400 mJ/cm^2^ of 365 nm UV light on ice. 4-SU can enhance the cross-linking efficiency especially for proteins in the cytoplasm [29, 30, 32]. The cells were centrifuged and the pellet stored at -80°C. The major steps of PAR-iCLIP are listed in Fig. S8 and detailed in the supporting information. CLIP experiments require an antibody, which can effectively immunoprecipitate tau under stringent high salt wash conditions. HJ 8.5 [62] raised against human tau efficiently depleted tau. With a dissociation constant of 0.3 pM, HJ 8.5 pulled down tau under the high-stringency purification conditions of CLIP and remained bound to tau throughout the procedure. Control experiments with GFP or CDK5 antibody for the lysate expressing those proteins, or with HJ 8.5 antibody for lysates without expressing tau were always done in parallel to rule out false positive binding caused by the beads. After immunoprecipitation, the cross-linked RNA was radiolabeled, the protein separated on an SDS-PAGE gel, and transferred to nitrocellulose membrane and blotted. The RNA-protein complexes from CLIP experiments (Fig. 1a-c) were cut from the blot, and the RNA extracted followed by reverse transcription for library preparation [30]. The ^32^P-labeled RNA band was run on a polyacrylamide Tris-Borate Urea (TBU) denaturing gel to demonstrate the confirm the size of the complex.

### Library preparation, deep sequencing and bioinformatics analysis of iCLIP

The iCLIP libraries contained an experimental and a random barcode, which allowed multiplexing and the removal of PCR duplicates. After the barcodes were introduced, sample and control from one set of experiments were mixed to remove batch-to-batch variation. Libraries were sequenced on an Ion Torrent Proton (Thermo Fisher). Fastx collapser from FASTX-Toolkit was used to collapse reads and filter replicates resulting from the PCR based on the random barcode. Reads were separated into samples by the barcodes at the 5` ends of reads. The reverse transcription primers sequenced at both ends of the reads were trimmed with Cutadapt [63]. After trimming the barcodes, reads with 18 bps or more were kept and counted as total unique reads, and aligned to the human genome (hg19) by Bowtie2 [64]. RseQC [65] was used to evaluate the quality of sequencing and mapping reads. Alignments with scores equal to or greater than ten were kept for downstream analysis. Reads from these RNA pools were clustered by their alignments, and Pyicos tools [66] were used to identify the significant clusters. Clusters with at least five reads were retained and considered to contain target sites for RNA-tau crosslinking. Gene models for RefSeq mRNAs, tRNAs, rRNAs were downloaded from the UCSC genome browser. miRNAs were downloaded from miRBase (release 20) and other categories of RNAs were download from Ensemble (release 73). Cross-link sites were identified as the termination site of the sequencing based on the iCLIP protocol [67]. Individual tRNA genes from the UCSC genome browser were predicted by using tRNAscan-SE v.1.23. The secondary structure for each tRNA was obtained from GtRNAdb (http://gtrnadb.ucsc.edu/).

### Deep sequencing and analyses of small RNA expression in HEK cells and hiPSC-derived neurons

Small RNAs were extracted from HEK cells and hiPSC-derived neurons using miRNA isolation kit (mirVana^TM^). Library preparation was adapted from the Ion RNA-seq v2 (Thermo Fisher) protocol. cDNAs with size range from 30-100 bp were selected and sequenced. Reads were aligned to both human genomes and tRNA sequences. When mapping reads to tRNA sequences, the Bowtie read aligner [68] protocol was used, in which a maximum of two mismatches were allowed. Reads aligned to tRNAs were counted and analyzed with a custom Perl scripts.

### Recombinant tau and tau fragments

Full length recombinant human 4R2N tau, N-terminal truncated, microtubule binding domain containing, K18 tau (residues 244-372) and Δtau187 (residues 255-441 with a His-tag at the N-terminus) were used for *in vitro* studies. Methods for expression and purification of all recombinant tau variants are detailed in the supplementary text. Two variants of Δtau187 were prepared via site-direct mutagenesis: Δtau187/322C contains a C291S mutation, leaving only one cysteine at site 322, and Δtau187G272C/S285C contains C291S, C322S, G272C and S285C mutations, leaving two cysteines at sites 272 and 285.

### RNA gel mobility shift assay

The gel shift assay was performed with recombinant full length 4R2N and K18 tau and chromatographically purified unacetylated yeast tRNA^Lys^ (tRNA Probes) in 100 mM sodium acetate buffer at pH 7.0. The molar concentration of tRNA^Lys^ was accurately re-measured with UV spectrophotometry after base-hydrolization to account for the hyperchromic effect from the secondary and tertiary structure of tRNA [36]. RNA43 was purchased in a kit (Pierce) with the sequence of 5’-CCUGGUUUUUAAGGAGUGUCGCCAGAGUGCCGCGAAUGAAAAA-3’. The hydrolyzed tRNA and RNA43 samples were then quantified with UV spectrophotometry at 260 nm using an extinction coefficient of 0.025 (μg/ml)^-1^cm^-1^. For the gel shift assay, protein was incubated with tRNA at 37 °C for 10 minutes in the presence of 0.5 mM EDTA, 0.5 mM MgCl_2,_ 2 U SUPERase• In™ RNase Inhibitor (Thermo Fisher), 0.01% IGEPAL CA-630 (Sigma-Aldrich), and then applied to a TBE 8% Polyacrylamide Gel (Thermo Fisher). After gel separation, tRNA was stained with SYBR Gold II (Thermo Fisher). For quantitative analysis, the fraction of free and bound tRNA was quantified in ImageJ2 (National Institute of Health [69]).

### Isothermal Titration Calorimetry (ITC) experiments

Full length tau or K18 tau were dialyzed overnight into a specified buffer for ITC (20 mM ammonium acetate, pH 7). tRNA (from Baker’s yeast, Sigma-Aldrich) was re-suspended in the ITC buffer and concentration determined using a Nanodrop 1000 (Thermo Scientific). Experiments were run on a Nano ITC (TA Instruments), in which 300 μM tRNA was titrated (5 μl injections) into a 1 ml protein solution of 30 μM tau. Data was analyzed using the NanoAnalyze v3.6 software (TA Instruments). After subtracting the heat generated by tRNA titration into an empty buffer, the experimental data was fitted well with an independent binding model, but not with a cooperative nor to an independent two-site model. Fitting to a cooperative or to an independent two-site model gave the statistical error being over 100% for the first Ka value. This could be due to the low enthalpy governing the binding event that is observed in the ITC data, and could possibly be resolved with increased concentrations of the reactants, here tau and RNA. However, solubility limitations likely preclude these experiments from being realistically achievable. Of note, the second Ka value produced by the two site model is very similar to the values obtained from an independent model, suggesting that the predominant interaction is indeed the one we described. Experiments were repeated in triplicates and standard error of the mean reported.

### Spin labeling of Δtau187

To achieve labeling with paramagnetic or diamagnetic probes, the protein was dissolved in 6 M guanidinium hydrochloride and labeled overnight at 4°C using a 20-fold molar excess of the spin label (1-oxyl-2,2,5,5-tetramethylpyrroline-3-methyl) methanethiosulfonate (MTSL, Toronto Research Chemicals) or the diamagnetic analog of MTSL (1-Acetoxy-2,2,5,5-tetramethyl-δ-3-pyrroline-3-methyl) methanethiosulfonate (Toronto Research Chemicals). Excess label was removed using a PD-10 desalting column (GE Healthcare) equilibrated in a 20 mM ammonium acetate buffer at pH 7.0. The protein was concentrated using a 3 kDa centrifugal filter (Amicon UFC800396). The final protein concentration was determined by UV-Vis absorption at 274 nm using an extinction coefficient of 2.8 cm^-1^mM^-1^, calculated from extinction coefficient of Tyrosine [70]. The two variants Δtau187/322C and Δtau187G272C/S285C were spin labeled with paramagnetic MTSL probes at the one or two cysteine sites, and are respectively referred to as Δtau187/322C-SL and Δtau187G272C/S285C-SL_2_. In order to achieve spin dilution, Δtau187/322C and Δtau187G272C/S285C were also labeled with the diamagnetic analogue of MTSL probes, and are referred to as Δtau187/322C/322C-DL and Δtau187G272C/S285C-DL_2_. Images shown in Fig. 4a-b were taken using 400 μM tau, 800 μg/ml poly(A) RNA and 30% glycerol in 20 mM ammonium acetate at pH 7.

### Preparation of tau-RNA complex coacervate

Droplets were formed in the 20 mM ammonium acetate buffer with NaCl concentration varying between 0 and 100 mM and glycerol concentration from 0 to 50% v/v. Unless explicitly indicated for measurements at various temperatures, all samples were prepared and characterized at room temperature. Solutions of Δtau187 or full length 4R2N tau was mixed with tRNA (Baker’s yeast, Sigma-Aldrich), poly(A) RNA or poly(U) RNA (Sigma-Aldrich) at varying protein, RNA, NaCl and glycerol concentrations in a 0.6 ml Eppendorf tube. RNAs were weighted out as powder and the mass concentration was calculated. The tau:RNA mass ratio, the total concentration of tau and RNA, as well as the NaCl salt concentration were optimized to maximize droplet formation, while choosing a total biopolymer density to avoid overlapping of droplets in the images to simplify the calculation of the droplet coverage (%). Microscopy images were acquired at 10 minutes after mixing—the droplets form spontaneously and are expected to be in equilibrium with the dilute supernatant. A concentration of 19% v/v for glycerol/water was determined to be an optimal concentration to ensure cryoprotection for DEER measurements carried out at ∼80 K, while also ensuring maximal droplet formation at room temperature (Fig. S6b).

### Bright-field microscopy of tau-RNA droplets

Immediately after mixing tau in a 0.6 ml Eppendorf tube with RNA under droplet forming conditions (established above) and ensuring thorough mixing, 1 μl of this mixture was applied to a microscope slide that is closed with a cover slide gapped by two layers of double-side sticky tape to generate a liquid sample region with consistent thickness. The microscope slide was kept at room temperature for 10 minutes with the cover slide facing down, during which the particles within the liquid sample region settled down onto the surface of the cover slide. Images were taken with a 12-bit CCD camera across the entire sample liquid region near the surface of the cover slide using an inverted compound microscope (Olympus IX70). Before imaging, Köhler illumination was applied and the focus close to the surface of the cover slide optimized to enhance the contrast between the dark droplets and the bright background.

### Confocal microscopy of tau-RNA droplets

For confocal microscopy, Δtau187/322C was fluorescence-labeled (Δtau187/322C-FL) with Alexa Fluor 488 C_5_ Maleimide (Thermo Fisher) at the same 322 site as Δtau187/322C-SL. 50 μM of Δtau187/322C-FL was mixed with 350 μM Δtau187/322C-SL at a 1:7 molar ratio in order to prevent saturation. 800 μg/ml of poly(A) RNA was further added to this tau solution, resulting in droplet formation in the 20 mM ammonium acetate buffer and in the presence of 20 mM NaCl. 10 μl of the mixture was pipetted and put on a microscope slide with a cover slide gapped by double-sided sticky tape. Confocal images were acquired using a spectral confocal microscope (Olympus Fluoview 1000).

### Droplet quantification from image analysis

To quantify the amount of droplet formed under the given experimental condition of interest, images were taken by a 12-bit CCD camera of an inverted compound microscope (Olympus IX70), and recorded in TIF format. With illumination and focus optimized, droplets settling on the cover slide have lower intensity than their surrounding on the images. An image of the 20 mM ammonium acetate buffer was taken to calculate the average intensity to set as threshold in order to classify different parts of the image into droplets and buffer. For each image, the area of the droplets was divided by the total area of the image, generating a % droplet coverage value on the cover slide. Droplets with eccentricity above 0.9 or equivalent diameter below 1μm were filtered out in order to reduce false reading. The MATLAB code is made available as supplementary files on the internet (https://github.com/yanxianUCSB/DropletAnalysis).

### Continuous wave (cw) Electron Paramagnetic Resonance (EPR)

Cw EPR relies on the anisotropy of nitroxide radical’s Larmor frequency and hyperfine coupling that makes its lineshape highly sensitive to the local dynamics, orientation and confinement of a nitroxide-based spin label tethered to the protein. Cw EPR measurements were carried out with Δtau187/322C-SL using a X-band spectrometer operating at 9.8 GHz (EMX, Bruker Biospin) and a dielectric cavity (ER 4123D, Bruker Biospin). Samples were prepared by either mixing 200 μM Δtau187/322C-SL with 300 μM unlabeled Δtau187/322C (to generate 40% spin labeled sample) or by using 500 μM Δtau187/322C-SL (100% labeled). Viscogen was added to the sample to achieve either 19% v/v glycerol (for the droplet samples) or 30% v/v sucrose (for the aggregated samples) matching the DEER conditions. tau samples under droplet forming condition were prepared by adding 1.5 mg/ml RNA, and tau samples under aggregation-inducing conditions prepared by adding 125 μM heparin (11 kDa average MW, Sigma-Aldrich). A sample of 3.5 μl volume was loaded into a quartz capillary (VitroCom, CV6084) and sealed at one end with critoseal and the other with beeswax, and then placed in the dielectric cavity for measurements. Cw EPR spectra were acquired using 2 mW of microwave power, 0.3 gauss modulation amplitude, 150 gauss sweep width and 25 scans for signal averaging.

### Double Electron Electron Resonance (DEER)

DEER measurements were performed on a Q-band pulsed EPR spectrometer operating at 32 GHz (E580, Bruker Biospin) equipped with a QT2 resonator (measurements done by courtesy of Bruker Biospin). Samples were prepared by mixing 50 μM Δtau187G272C/S285C-SL_2_ with 500 μM analog-labeled Δtau187G272C/S285C-DL_2_ at a 1:10 molar ratio to achieve spin-dilution and avoid artifacts from unwanted inter-protein spin distances. For DEER, tau samples under droplet forming condition were prepared by adding 1.65 mg/ml RNA and ensuring 19% v/v glycerol concentration, and tau samples under aggregation-inducing conditions prepared by adding 137.5 μM heparin and ensuring 30% v/v sucrose concentration. 40 μL samples containing 550 μM concentration of tau were loaded into a quartz capillary (2.4 mm od x 2 mm id) and flash frozen in liquid nitrogen after 20 minutes of incubation at room temperature and the specific conditions listed for the data. DEER measurements were conducted using the dead-time free four-pulse DEER sequence at 80 K, using 22 ns (π/2) and 44 ns (π) observe pulses and a 30 ns (π) pump pulse. The raw DEER data was processed using Gaussian fitting via DeerAnalysis2013 [71].

### Quantification of fibril using Thioflavin T assay

160 μM Δtau187 was mixed with 480 μg/ml poly(U) RNA or 40 μM heparin and incubated at room temperature in the presence of 4 μM Thioflavin T for over 15 h (Fig. S6f). The fluorescence intensity at 485 nm was measured using a plate reader (Tecan Infinite 200 Pro).

### Transfer RNA transfection

Transfection of RNA in neuron was first validated using tRNA^Phe^-fluorescein transfected in primary mouse neuron at 14 days *in vitro* using lipofectamine 2000 transfection reagent (Thermo Fisher) following the manufacturer’s protocol (Fig. S7) Cell nuclei was stained with DAPI before fluorescence microscopic imaging.. For sarkosyl experiments hiPSC neuronal cultures containing wild type or mutant tau were transfected with 48 μg of bovine liver tRNA (Sigma-Aldrich) per six well plate. Control cells were transfected in equal conditions in the absence of nucleic acid (Mock transfection). Mock transfected cells were lysed as described below, with or without 48 μg tRNA added to the lysis buffer.

### Sarkosyl insolube tau isolation and western blotting

Separation of sarkosyl insoluble tau was done as described in the literature [72, 73]. Briefly, adherent neuronal cell cultures were lysed with an ice-cold high salt/high sucrose Tris HCl buffer (0.8 M NaCl, 10% Sucrose, 10 mM Tris HCl pH 7.4) containing a 1X Protease Inhibitor Cocktail, and a 1X Phosphatase Inhibitor Cocktail. Lysis proceeded at 4°C for 30 min before detachment with a cell scraper, followed by mechanical dissociation using a micropipette. Immediately afterwards, the lysates were centrifuged at 4°C 3000 x g for 15 min in a microcentrifuge. The clear supernatants were collected and sampled (Input). Sodium lauroyl sarcosinate (Sigma-Aldrich) was then added to the supernatants to a final concentration of 1%, followed by a brief vortexing. Samples were incubated at 4°C with continuous rocking. Samples were then centrifuged at 4°C for 2 h at 170000 x g in a Beckman Coulter 70.1 Ti rotor. Sarkosyl soluble supernatants (Sol.) were collected. Sarkosyl insoluble (Ins.) pellets were resuspended in 2X sample loading buffer (250 mM Tris HCl pH 6.8, 10% Glycerol, 10% SDS, 0.5% bromophenol blue, 20 mM DTT) and heated to 95°C for 10 min. The previously collected input and the sarkosyl soluble fractions were diluted with the same sample loading buffer, and heated under similar conditions. Proteins were separated on a 10% SDS-PAGE, transferred to Nitrocellulose membranes, and blotted with either PHF-1 or β–actin antibody. Western blot signal was detected with a LI-COR Odyssey imaging system (LI-COR Biosciences) and quantified in ImageJ2 (National Institute of Health [69].

### Data availability

The data that support the findings of this study are available from the corresponding authors upon request.

## Acknowledgments

We are grateful to the tau consortium for financial support to K.S.K. and S.H. Thanks to D. M. Holtzman, Washington University School of Medicine for the generous gift of HJ 8.5 and HJ 9.2 antibodies and P. Davies, Albert Einstein College of Medicine for PHF-1 antibody. This work was supported by the NIH grant (#R01AG05605)) to S.H and K.S.K, the 2011 NIH Director New Innovator Award to S.H and made use of the Material Research Laboratory (MRL) Central Facilities supported by the National Science Foundation (NSF) through the Materials Research Science and Engineering Center under Grant DMR 1121053. We acknowledge the use of the NRI-MCDB Microscopy Facility and the Spectral Laser Scanning Confocal supported by the Office the Director, National Institutes of Health (NIH) under Award # S10OD010610. The authors declare no competing financial interests.

